# Reduced occurrence of alpha waves during resting state predicts high ADHD traits in young adults

**DOI:** 10.1101/2025.04.11.648375

**Authors:** Julio Rodriguez-Larios, Ümit Aydin, Gráinne McLoughlin

## Abstract

Attention-deficit/hyperactivity disorder (ADHD) is a neurodevelopmental condition with significant cognitive and social impacts. Identifying reliable biomarkers for ADHD is crucial for developing personalized therapies. Electroencephalography (EEG) alpha oscillations (8–12 Hz) have been suggested as a potential biomarker, but findings have been inconsistent. This study aimed to investigate whether alpha oscillations in young adulthood are associated with high ADHD traits using EEG data from a large twin sample (N = 556) enriched with participants with ADHD and autism. We assessed whether alpha oscillations during rest were associated with high ADHD traits through logistic regression analysis. In addition, we used twin modelling to estimate the heritability of EEG alpha measures and their relationship with ADHD traits. Results showed that relative alpha power was a significant predictor of ADHD traits when controlling for other factors such as age, sex and autistic traits. Specifically, we found that for each unit decrease in relative alpha power, the likelihood of being in the high ADHD trait group increased by approximately 26%. Further analysis suggested that group differences were due to a reduced occurrence (but not amplitude) of oscillatory bursts in the alpha range. Finally, our twin modelling results suggested that although alpha power is heritable, the genetic factors contributing to individual differences in alpha measures and ADHD traits were largely independent. Together, these findings suggest that reduced alpha oscillations, particularly occurrence of alpha bursts, may serve as a potential biomarker for ADHD. Our results may have implications for neuromodulation therapies targeting alpha rhythms in ADHD, such as neurofeedback and transcranial alternating current stimulation (tACS).

## Introduction

Attention-deficit/hyperactivity disorder (ADHD) is a neurodevelopmental condition characterized by pervasive patterns of inattention, hyperactivity, and impulsivity that is estimated to affect approximately 5% of adults worldwide (McCarthy et al., 2012; Polanczyk et al., 2007). ADHD has a significant impact on individuals’ quality of life, mental health and social relationships (Adamou et al., 2013; Aduen et al., 2018; Aydin et al., 2022a; Capp et al., 2023). While psychopharmacological treatments, particularly stimulant medications, are considered the most effective interventions for managing ADHD symptoms (Cortese et al., 2018; Crescenzo et al., 2017; Skirrow et al., 2014), their use is frequently accompanied by side effects such as loss of appetite, sleep problems, and mood disturbances (Cascade et al., 2010). Although non-pharmacological treatments show high promise, the evidence of their success is still limited (Nimmo-Smith et al., 2020).

Recent advancements in precision psychiatry and neuromodulation have paved the way for developing personalized therapies for ADHD (Arns, 2012; Voetterl et al., 2023). In this regard, neuromodulation techniques, such as neurofeedback and non-invasive brain stimulation, seem promising alternative or complementary treatments (Arns et al., 2014; Boetzel & Herrmann, 2021; Frohlich & Riddle, 2021). These approaches aim to directly modulate brain activity to improve symptomatology by promoting neuroplastic changes (Arns et al., 2014; Veniero et al., 2015; Vossen et al., 2015). However, a critical prerequisite for the success of neuromodulation-based therapies is the identification of reliable brain biomarkers that can guide intervention strategies (Klooster et al., 2024; Riddle & Frohlich, 2023). Without clear and replicable biomarkers, the development and optimization of neuromodulation techniques remain limited.

The study of brain oscillations in ADHD has emerged as a particularly promising approach for identifying biomarkers relevant for neuromodulation (Günther et al., 2022; McLoughlin, Makeig, et al., 2014; Riddle & Frohlich, 2021). Neural oscillations, as measured by electroencephalography (EEG), reflects the coordinated firing of neuronal populations and has been implicated in various cognitive and behaviouralprocesses (Klimesch, 1999; von Stein & Sarnthein, 2000). The study of oscillatory activity in ADHD has been dominated by investigations of the theta-beta ratio (a higher ratio between theta (4-7 Hz) and beta (15-30 Hz) oscillations) (Arns et al., 2013; McLoughlin et al., 2022). However, recent evidence has challenged the reliability and specificity of this finding (for a review, see McLoughlin et al., 2022). Alpha oscillations (8–12 Hz) have emerged as a frequency band of interest in ADHD research, with several studies reporting to be reduced in individuals with ADHD (Cañigueral et al., 2022; Deiber et al., 2020; Loo et al., 2009; Voetterl et al., 2023; Woltering et al., 2012). However, the literature remains inconsistent with other studies reporting no significant differences (Bender et al., 2023; Bresnahan et al., 2006; Liu et al., 2016). These discrepancies may arise from small sample sizes or confounding factors such as age, sex and co-occurring conditions(Brandeis et al., 2018; Chen et al., 2024; Loo et al., 2018; McLoughlin et al., 2022). Another consideration is the influence of non-oscillatory components of the EEG signal, which may confound traditional measures of band-specific power (Chen et al., 2024; Donoghue et al., 2021; Robertson et al., 2019).

In the present study, we investigated whether alpha oscillations are associated with high ADHD traits in a large EEG sample of twins (N = 556). Specifically, we compared relative alpha power between individuals with high (N = 193) and low ADHD traits (N = 343) using a logistic regression analysis that controlled for covariates including age, sex and autistic traits. In addition, we assess whether putative differences in spectral power are actually due to oscillatory activity using a recently developed algorithm that detects oscillatory bursts in EEG (Rodriguez-Larios et al., 2024; Rodriguez-Larios & Haegens, 2023). Finally, we used twin modelling to assess the genetic and environmental contributions to variance in alpha oscillatory dynamics, allowing us to estimate the heritability of the relevant alpha power measures and their relationship with ADHD traits.

## Methods

### Participants

Ethical approval for the study was received from King’s College London Psychiatry, Nursing and Midwifery Research Ethics Subcommittee (RESCMR-16/17-2673). Data were collected as part of the Individual Differences in EEG in young Adults Study (IDEAS) which is a subsample of the Twins Early Development Study (TEDS) (Rimfeld et al., 2019). The sample consisted of 556 participants (267 males) with an average age of 22.43 (SD = 0.96 years) (119 Monozygotic (MZ) and 164 Dizygotic (DZ) pairs) (see full sample description at (Aydin et al., 2022a)).

Barkley Adult ADHD rating scale-IV (BAARS-IV) and Diagnostic interview for ADHD in adults 2.0 (DIVA 2.0) were used to identify participants with high and low ADHD traits. Participants were classified as having high ADHD traits if they met DSM-5 diagnostic criteria for adult ADHD based on self-reported data from the DIVA 2.0 or if their BAARS-IV total ADHD score was 39 or higher, reflecting mild to significant ADHD traits and related difficulties (Capp et al., 2023). This resulted in 205 participants in the high ADHD traits group and 351 in the low ADHD traits group.

In order to control for autistic traits, Autism Diagnostic Observation schedule 2 (ADOS-2) and Social Responsiveness Scale-2 (SRS-2) were used to classify participants into those with high (N = 127) and low (N = 429) autistic traits. Individuals with high autistic traits were the ones that scored four or higher on the ADOS-2 CSS (indicating behaviours consistent with an autism diagnosis) or had an SRS-2 raw score of 68 or above (indicating mild to severe difficulties related to autistic social traits). Those who did not meet these criteria were classified as having low autistic traits (Capp et al., 2023).

### EEG recording and analysis

EEG was recorded with a mobile wireless 64-channel (10-10 montage) system (Cognionics, San Diego, CA; Ag/AgCl electrodes, with a sampling rate of 500 Hz). The dataset contained both resting state (eyes open and eyes closed) and task EEG data (Aydin, Gyurkovics, et al., 2023). Three minutes long eyes closed resting state EEG data was used in this study. This choice was based on previous literature showing higher reproducibility of eyes closed resting state EEG (Duan et al., 2021).

EEG analysis was performed in MATLAB R2023a using custom scripts and EEGLAB (Delorme & Makeig, 2004). Data cleaning was performed using an automatic pre-processing pipeline based on EEGLAB functions. First, data were resampled to 250 Hz, re-referenced to common average and filtered between 1 and 30 Hz. Abrupt noise in the data was removed using the Artifact Subspace Reconstruction method with a cut-off value of 20 SD (Chang et al., 2019). Noisy channels were detected automatically with the EEGlab function *clean_channels* using a threshold of 0.6 and were later interpolated. The average number of interpolated channels was 9.5. Independent component analysis (ICA) and an automatic component rejection algorithm (Pion-Tonachini et al., 2019) were used to discard components associated with muscle activity, eye movements, heart activity or channel noise (threshold = 0.8). The average number of rejected components was 2.5. Continuous data was divided into 2 second epochs (50 % overlap) and epochs that exceeded an amplitude of ±100 mV were removed (mean number of rejected epochs = 18). Subjects with more than 50 % of rejected epochs were excluded from subsequent analysis (N = 20).

For the estimation of relative alpha power, the power spectrum between 1 and 30 Hz (in steps of 0.5 Hz) was estimated using a short-term Fourier transform as implemented through the MATLAB function *spectrogram* (sliding window of 2 seconds with 50 % overlap). Individual alpha peak was estimated as the local maximum between 7 and 14 Hz using the *findpeaks* MATLAB function. Subjects that did not show an alpha peak in a specific electrode were excluded from the statistical analysis. In this regard, 23 subjects did not show an alpha peak in at least one electrode. Relative alpha power was defined as the amplitude of the individual alpha peak of the normalised spectrum (i.e. z-scored over the frequency dimension).

In order to assess whether putative differences in relative alpha power were due to oscillatory activity, a recently developed algorithm was adapted (Rodriguez-Larios et al., 2024; Rodriguez-Larios & Haegens, 2023). In short, EEG data is first transformed to the time-frequency domain using Morlet wavelets (6 cycles width) as implemented in (Caplan et al., 2015). The frequency resolution was 1Hz (between 1 and 30 Hz). Oscillatory bursts are defined as time points in which the amplitude at a specific frequency exceeds estimate of aperiodic activity (e.g. 1/f trend) for at least one full cycle. The 1/f trend is estimated by fitting a straight line to the spectrum in the log-log space per electrode across all time points and epochs. Only oscillatory bursts that had their highest spectral peak in the alpha range (7-14 Hz) were selected. This algorithm allows to disentangle the quantity of time in which oscillatory activity is present (i.e. oscillatory burst coverage) and the amplitude of the burst (which is estimated as the oscillation maximum peak prominence in the time domain). MATLAB code of the of the entire analysis pipeline can be found here: https://osf.io/yef9z/?view_only=c14d995a2b464be68fdfad4cc11f56a7

### Statistical analysis

A logistic regression model was employed to investigate the association between EEG alpha oscillations and the likelihood of having high ADHD traits. The analysis was conducted using a generalized linear mixed-effects model (GLMM) with a logit link function in MATLAB 2023a (*fitglme* function). To control for potential confounding factors, the model included additional fixed effects for age, sex and autistic traits (binary: high vs low). To account for clustering within the sample due to relatedness (i.e. twins), random intercepts were included in the model for each pair of twins (Aydin, Cañigueral, et al., 2023; Malone et al., 2014). Only the association between alpha oscillation variables and ADHD traits will be discussed as the rest of the predictors were only included as controls. The regression coefficients (β) were exponentiated to calculate the odds ratio. In order to assess significance, t-values for fixed effects are calculated as the ratio of the estimated coefficient to its standard error.

Correlations were computed through Pearson correlation coefficient and multiple comparisons correction was implemented through the False Discovery Rate (Benjamini & Hochberg, 1995). Outliers were detected and rejected before all statistical analysis. Outliers were defined as elements more than three scaled median absolute deviations from the median (see *isoutlier* function in MATLAB 2023a).

### Twin modelling

We applied genetic multivariate liability threshold models based on the principles that monozygotic (MZ) twins share 100% of their genetic influences, while dizygotic (DZ) twins share 50%, and both groups share common environmental factors equally (Neale & Maes, 2004). These models treat high ADHD traits and high autistic traits as dichotomous variables and use the MZ:DZ ratio of cross-twin within-trait correlations to partition the variation in EEG traits into standardised additive genetic variance (a²), common environmental variance (c²), and unique environmental variance, including measurement error (e²). The phenotypic correlation (Rph) between traits is calculated as the zero-order correlation, while the MZ:DZ ratio of cross-twin cross-trait correlations is used to partition the covariation between traits (e.g., ADHD and relative alpha power) to report genetic (Ra) correlations. Since the sample had a higher proportion of participants meeting ADHD or ASD criteria compared to the general population, we applied corrections to the standard twin model, as described in (Aydin et al., 2022b). Twin modelling was conducted using structural equation modelling in OpenMx (Boker et al., 2011), with likelihood-based asymmetric 95% confidence intervals (CIs) estimated for all parameters. To examine associations between ADHD or autistic traits and specific EEG metrics, we ran bivariate models for relative alpha power, burst alpha amplitude, and burst alpha coverage.

## Results

### Differences between groups in relative alpha power

Our results showed that relative alpha power was a significant predictor of ADHD traits (p-values after FDR correction < 0.05 in several electrodes) (see **Figure 1A**). Specifically, participants with high ADHD traits showed a significant reduction in relative alpha power when compared to the low ADHD trait group (see **Figure 1B-C**). Although the effect was significant in the majority of electrodes, it was more pronounced in frontal electrodes (see the most significant electrode (AF8) marked in **Figure 1A**). The odds ratio of this electrode was 0.74, which means that for every unit of relative alpha power (in z-scores) the odds of being in the high ADHD trait group decreased by approximately 26 % (t-value (497) = −3.02; p-value = 0.0026). The complete regression model is summarised in **Table 1** for this electrode.

**Figure 1.**
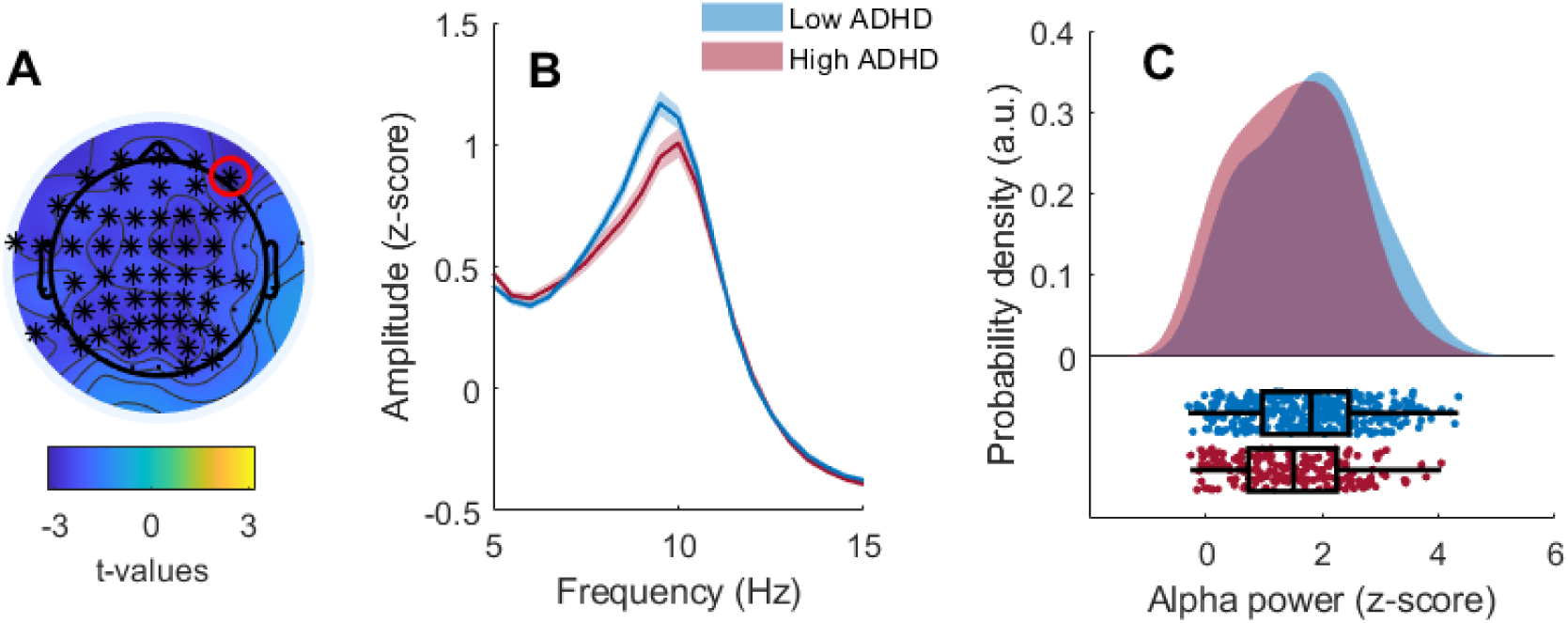
Association between relative alpha power and ADHD traits. A) Topographical plot depicting t-values obtained when using relative alpha power as predictor in the specified model. Asterisks indicate significant electrodes (*p-value* <0.05) after FDR correction and the red circle marks AF8, the most significant electrode (i.e. highest absolute t-value). B) Power spectrum (z-scored) averaged across significant electrodes for low (blue) and high (red) ADHD trait groups. The shaded area depicts standard error. C) Relative alpha power for the low (blue) and high (red) ADHD trait groups. Each point depicts relative alpha power for one subject across significant electrodes. Boxplots indicate the median and interquartile range per group.

**Table 1.** Logistic regression model for the prediction of ADHD traits (0 = low,1 = high) from autistic traits (0 = low,1 = high), age, sex and relative alpha power at electrode AF8.

| Predictor | $\beta$ | SE | t-value | p-value | Odds ratio |
| --- | --- | --- | --- | --- | --- |
| *Intercept* | 6.18 | 2.59 | 2.38 | 0.0172 | 487.23 |
| *Autistic traits* | 1.02 | 0.23 | 4.28 | <0.001 | 2.78 |
| *Age* | -0.29 | 0.11 | -2.63 | 0.0087 | 0.74 |
| *Sex* | 0.03 | 0.21 | 0.17 | 0.8610 | 1.04 |
| *Relative alpha power* | -0.323 | 0.10 | -3.02 | 0.0026 | 0.72 |

### Assessing the putative oscillatory nature of the group differences

In order to assess whether the reported differences in alpha power were due to oscillatory activity, we employed a recently developed algorithm that estimates the occurrence and amplitude of oscillatory bursts (Rodriguez-Larios et al., 2024; Rodriguez-Larios & Haegens, 2023). For this analysis, we selected the frontal EEG electrode with the highest group difference as quantified by their t-value (i.e. electrode AF8; see red circle in **Figure 1**).

Logistic regression analysis revealed a significant effect of oscillatory burst coverage (t-value (499) = - 2.28, p-value = 0.0228) but not of oscillatory burst amplitude (t-value (499) = 0.13, p-value = 0.89) (see **Table 2** for full model details and **Figure 2A-B** for depiction of results). The odds ratio for burst coverage was 0.94, which means that for every second of alpha oscillatory bursts in the resting state EEG, the odds of being in the high ADHD trait group decreased by approximately 6 %. These results suggest that relative alpha power differences between groups was due to the occurrence of oscillatory bursts rather than their amplitude. In line with this idea, inter-individual differences in relative alpha power showed a higher correlation to burst coverage (*r-value* = 0.83; *p-value* <0.001; **Figure 2C**) than to burst amplitude (*r-value* = 0.33; *p-value* <0.001; **Figure 2D**).

**Figure 2.**
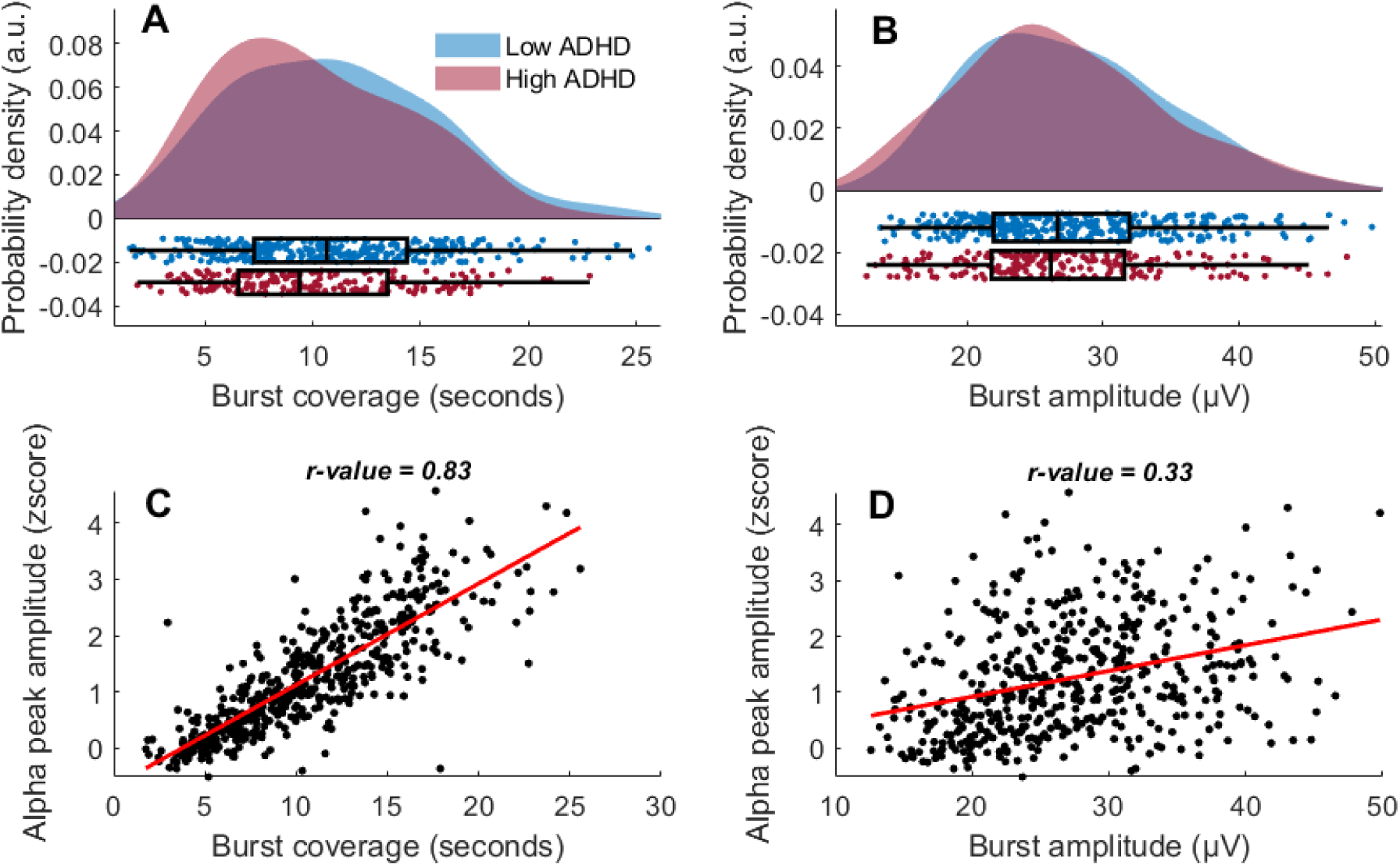
Alpha oscillatory bursts differences between groups with high and low ADHD traits and their correlation with relative alpha power. A) Burst coverage for the low (blue) and high (red) ADHD trait groups. Each point depicts relative alpha power for one subject for the selected frontal electrode. Boxplots indicate median and interquartile range per group. B) Same as A but for burst amplitude. C-D) Correlation between burst parameters (coverage and amplitude) and individual alpha peak amplitude (each point depicts data of one subject for the selected electrode).

**Table 2.** Logistic regression model for the prediction of ADHD trait (0 = low,1 = high) from autistic trait (0 = low,1 = high), age, sex, mean oscillatory burst alpha amplitude (i.e. Amplitude) and mean oscillatory burst alpha coverage (Coverage) at electrode AF8.

| Predictor | $\beta$ | SE | t-value | p-value | Odds ratio |
| --- | --- | --- | --- | --- | --- |
| *Intercept* | 4.90 | 2.46 | 1.98 | 0.0472 | 134.93 |
| *Autistic traits* | 1.04 | 0.23 | 4.48 | <0.001 | 2.84 |
| *Age* | -0.23 | 0.10 | -2.15 | 0.0316 | 0.79 |
| *Sex* | 0.04 | 0.20 | 0.22 | 0.8245 | 1.04 |
| *Amplitude* | 0.002 | 0.01 | 0.13 | 0.8901 | 1.00 |
| *Coverage* | -0.06 | 0.02 | -2.28 | 0.0228 | 0.94 |

### Assessing the genetic contributions

We found statistically significant and high genetic contributions (a^2^) to the total variance for relative alpha power and burst alpha coverage but not for burst alpha amplitude (**Table 3**). Common environment contributions (c^2^) were not significant.

**Table 3:** Phenotypic (Rph) and genetic (Ra) correlations between ADHD trait and relative alpha power, mean oscillatory burst alpha amplitude, and **mean oscillatory burst alpha coverage.** Standardised estimates of genetic (**a^2^**), common (**c^2^**) and nonshared (**e^2^**) environment contributions to variance of the EEG measures. 95% confidence intervals are given in parentheses, and significant estimates are written in bold.

|  | Rph | Ra | a<sup>2</sup> | c<sup>2</sup> | e<sup>2</sup> |
| --- | --- | --- | --- | --- | --- |
| **Relative alpha power** | **-0.15**<br>(-0.25,-0.05) | -0.14<br>(-0.31,0.03) | **0.59**<br>(0.48,0.69) | 0.00<br>(0.00,0.28) | **0.41**<br>(0.31,0.52) |
| **Burst alpha amplitude** | -0.04<br>(-0.14,0.05) | -0.05<br>(-1.00,1.00) | 0.38<br>(0.00,0.61) | 0.11<br>(0.00,0.44) | **0.51**<br>(0.39,0.65) |
| **Burst alpha coverage** | **-0.12**<br>(-0.22,-0.03) | -0.13<br>(-0.31,0.05) | **0.59**<br>(0.46,0.69) | 0.00<br>(0.00,0.29) | **0.41**<br>(0.31,0.54) |

In terms of associations between ADHD traits and alpha measures, we observed significant negative phenotypic correlations for relative alpha power (−0.15) and burst alpha coverage (−0.12) but in both cases, the genetic correlations (Ra) were not significant (**Table 3**). Phenotypic and genetic correlations between autistic traits and the alpha measures were not statistically significant (see **Table 4**).

**Table 4:** Phenotypic (Rph) and genetic (Ra) correlations between autistic trait and relative alpha power, mean oscillatory burst alpha amplitude, and **mean oscillatory burst alpha coverage.** Standardised estimates of genetic (**a^2^**), common (**c^2^**) and nonshared (**e^2^**) environment contributions to variance of the EEG measures. 95% confidence intervals are given in parentheses, and significant estimates are written in bold.

|  | Rph | Ra | a<sup>2</sup> | c<sup>2</sup> | e<sup>2</sup> |
| --- | --- | --- | --- | --- | --- |
| **Relative alpha power** | -0.07<br>(-0.19, 0.05) | -0.09<br>(-0.30,0.12) | **0.59**<br>(0.26,0.69) | 0.00<br>(0.00,0.28) | **0.41**<br>(0.31,0.52) |
| **Burst alpha amplitude** | -0.08<br>(-0.05,0.20) | 0.27<br>(-0.02,1.00) | 0.35<br>(0.00,0.61) | 0.13<br>(0.00,0.44) | **0.51**<br>(0.39,0.65) |
| **Burst alpha coverage** | -0.11<br>(-0.23,0.01) | -0.20<br>(-0.41,0.02) | **0.59**<br>(0.46,0.69) | 0.00<br>(0.00,0.27) | **0.41**<br>(0.31,0.54) |

## Discussion

The aim of this study was to investigate whether adults with high ADHD traits show alterations in alpha oscillations during resting state. For this purpose, we analysed a large twin EEG data set (N = 556) enriched for ADHD and autistic traits (Aydin et al., 2022a; McLoughlin, Palmer, et al., 2014). Our results showed that participants with high ADHD traits have significantly reduced relative alpha power at rest across EEG electrodes (although effects were more pronounced in frontal electrodes). Specifically, logistic regression analysis revealed that for every unit of relative alpha power (in z-scores), the odds of being in the high ADHD trait group decreased by approximately 26%. Further analysis suggested that group differences in relative alpha power were due to changes in the occurrence of oscillatory bursts in the alpha range rather than their amplitude. Finally, our twin modelling analysis confirmed the high heritability of alpha, specifically the occurrence of oscillatory bursts.

Our study is in line with previous literature showing that relative alpha power during resting state is significantly reduced in populations with high ADHD traits (Deiber et al., 2020; Loo et al., 2009; Ponomarev et al., 2014; Woltering et al., 2012). However, this finding is not completely consistent in the literature since other studies found no differences or even significant increases in resting state alpha power in adults with high ADHD traits (Bresnahan et al., 2006; Koehler et al., 2009; Poil et al., 2014; van Dongen-Boomsma et al., 2010; Voetterl et al., 2023). Our large well-characterised sample allowed us to attempt to clarify these inconsistencies by demonstrating that reduced alpha power remains significantly associated with ADHD even when controlling for autistic traits, age and sex. Another possible confounder in previous investigations of alpha and ADHD could be differences in the analytical approach. In this way, the majority of previous studies quantified alpha oscillations without controlling for aperiodic (non-oscillatory) activity, which has been recently shown to be affected in ADHD (Dakwar-Kawar et al., 2024). We employed a recently developed algorithm (Rodriguez-Larios et al., 2024; Rodriguez-Larios & Haegens, 2023) to investigate the origins of the reported alpha power differences in relation to ADHD traits. Crucially, this algorithm not only controls for the influence of aperiodic activity (Donoghue et al., 2021) but also allows for the distinction between the occurrence and amplitude of transient bursts of oscillatory activity. Using this algorithm, we demonstrated that alpha oscillations in the high ADHD trait group showed reduced occurrence (but not amplitude). In addition, interindividual differences in relative alpha power had a higher correlation to oscillatory burst occurrence than to burst amplitude. This analysis suggests that alpha power changes can be more influenced by the occurrence of alpha bursts than their amplitude. Distinguishing between modulations in amplitude and occurrence of oscillatory bursts can be crucial from a translational perspective, since some neuromodulation protocols have been shown to alter the occurrence (but not the amplitude) of alpha oscillations (Ossadtchi et al., 2017).

Our twin modelling results confirmed the high heritability of EEG alpha measures, with particularly strong genetic contributions to relative alpha power and burst alpha coverage. This finding aligns with previous reports indicating a substantial genetic contribution to alpha measures (Malone et al., 2014; Smit et al., 2006; van Beijsterveldt & van Baal, 2002). Heritability refers to the proportion of variance in a trait that can be attributed to genetic differences among individuals. In this context, it indicates that individual differences in alpha power and burst coverage are predominantly driven by genetic factors, rather than environmental influences. Despite the observed phenotypic correlations between ADHD traits and alpha oscillations, we did not detect significant genetic correlations between these measures. This suggests that while both alpha power and ADHD traits are heritable, the genetic factors contributing to each may be largely independent, and shared genetic factors are not critical for understanding the relationship between alpha oscillations and ADHD traits. While we observed a significant contribution of autistic traits in our logistic regression model (odds ratio=2.78), we did not observe significant phenotypic overlap in our twin model, which may indicate some specificity to ADHD.

The observed decrease in the occurrence of alpha oscillations in individuals with high ADHD traits could be due to several factors. One possibility is that these differences stem from group differences in brain anatomy, as alpha oscillations during rest have been linked to inter-individual differences in anatomical architecture (Kumral et al., 2022; Valdés-Hernández et al., 2010). In support for this idea, structural abnormalities in the prefrontal and anterior cingulate cortex have been reported in ADHD (Bayard et al., 2020). Another possibility is that alpha reductions reflect variability in resting-state cognition, as ADHD is associated with altered arousal and mind-wandering (Biederman et al., 2019; Lanier et al., 2021; Seli et al., 2015), which are known to highly influence alpha dynamics (Compton et al., 2019; Rodriguez-Larios et al., 2021; Rodriguez-Larios & Alaerts, 2020). Hence, in order to disentangle the origin of alpha reductions in ADHD, future research would need to incorporate both anatomical measures (e.g., individual MRIs) and resting state cognition questionnaires (see Diaz et al., 2013).

The identification of alpha oscillations as a possible biomarker for ADHD has important implications for therapy. If reduced alpha oscillations are present in ADHD, neuromodulation protocols that enhance alpha oscillations (such as transcranial alternating current stimulation and neurofeedback) (Soriano et al., 2024; Vossen et al., 2015) may improve ADHD symptoms. Recent evidence suggests that specific alpha oscillatory patterns may also help guide treatment selection, potentially improving clinical outcomes through personalised approaches (Voetterl et al., 2023). In this direction, it has been shown that a neurofeedback protocol that increased alpha power improved ADHD symptoms and attentional performance (Deiber et al., 2020). Similarly, a recent alpha tACS study has shown that repeated sessions of alpha tACS can improve ADHD symptoms (although in this case changes in EEG were not investigated) (Farokhzadi et al., 2020). Together, although more evidence is needed, neuromodulation protocols aimed at enhancing alpha oscillations in ADHD show significant promise. An important question for future research is whether individuals with ADHD who exhibit reduced alpha power at rest would derive greater benefits from these therapies.

In conclusion, this study shows that adults with high ADHD traits have reduced relative alpha power at rest, primarily due to a decrease in the occurrence of alpha oscillatory bursts. While our findings support the idea that alpha oscillations could be a potential biomarker for ADHD, its consistency across different populations and usefulness for therapy is still to be tested. Future research should investigate the underlying mechanisms contributing to these differences and determine if individuals with lower alpha power benefit more from targeted neuromodulation therapies.

## Acknowledgments

IDEAS was supported by UK Medical Research Council New Investigator Research grant (Grant No. MR/N013182/1 [to GM].

We gratefully acknowledge the participating families, and all staff involved in this study, in particular the IDEAS research team.

## Declaration of interests

The authors report no conflicts of interest.

## Notes

### Competing Interest Statement

The authors have declared no competing interest.

## References

Adamou, M., Arif, M., Asherson, P., Aw, T.-C., Bolea, B., Coghill, D., Guðjónsson, G., Halmøy, A., Hodgkins, P., Müller, U., Pitts, M., Trakoli, A., Williams, N., & Young, S. (2013). Occupational issues of adults with ADHD. BMC Psychiatry, 13(1), 59. 10.1186/1471-244X-13-59

Aduen, P. A., Day, T. N., Kofler, M. J., Harmon, S. L., Wells, E. L., & Sarver, D. E. (2018). Social Problems in ADHD: Is it a Skills Acquisition or Performance Problem? Journal of Psychopathology and Behavioral Assessment, 40(3), 440–451. 10.1007/s10862-018-9649-7

Arns, M. (2012). EEG-Based Personalized Medicine in ADHD: Individual Alpha Peak Frequency as an Endophenotype Associated with Nonresponse. Journal of Neurotherapy, 16(2), 123–141. 10.1080/10874208.2012.677664

Arns, M., Conners, C. K., & Kraemer, H. C. (2013). A Decade of EEG Theta/Beta Ratio Research in ADHD: A Meta-Analysis. Journal of Attention Disorders, 17(5), 374–383. 10.1177/1087054712460087

Arns, M., Heinrich, H., & Strehl, U. (2014). Evaluation of neurofeedback in ADHD: The long and winding road. Biological Psychology, 95, 108–115. 10.1016/j.biopsycho.2013.11.013

Aydin, Ü., Cañigueral, R., Tye, C., & McLoughlin, G. (2023). Face processing in young adults with autism and ADHD: An event related potentials study. Frontiers in Psychiatry, 14. https://www.frontiersin.org/articles/10.3389/fpsyt.2023.1080681

Aydin, Ü., Capp, S. J., Tye, C., Colvert, E., Lau-Zhu, A., Rijsdijk, F., Palmer, J., & McLoughlin, G. (2022a). Quality of life, functional impairment and continuous performance task event-related potentials (ERPs) in young adults with ADHD and autism: A twin study. JCPP Advances, 2(3), e12090. 10.1002/jcv2.12090

Aydin, Ü., Capp, S. J., Tye, C., Colvert, E., Lau-Zhu, A., Rijsdijk, F., Palmer, J., & McLoughlin, G. (2022b). Quality of life, functional impairment and continuous performance task event-related potentials (ERPs) in young adults with ADHD and autism: A twin study. JCPP Advances, 2(3), e12090. 10.1002/jcv2.12090

Aydin, Ü., Gyurkovics, M., Ginestet, C., Capp, S., Greven, C. U., Palmer, J., & McLoughlin, G. (2023). Genetic Overlap Between Midfrontal Theta Signals and Attention-Deficit/Hyperactivity Disorder and Autism Spectrum Disorder in a Longitudinal Twin Cohort. Biological Psychiatry, 94(10), 823–832. 10.1016/j.biopsych.2023.05.006

Aydin, Ü., Vorwerk, J., Küpper, P., Heers, M., Kugel, H., Galka, A., Hamid, L., Wellmer, J., Kellinghaus, C., Rampp, S., & Wolters, C. H. (2014). Combining EEG and MEG for the reconstruction of epileptic activity using a calibrated realistic volume conductor model. PLoS One, 9(3), e93154. 10.1371/journal.pone.0093154

Bayard, F., Nymberg Thunell, C., Abé, C., Almeida, R., Banaschewski, T., Barker, G., Bokde, A. L. W., Bromberg, U., Büchel, C., Quinlan, E. B., Desrivières, S., Flor, H., Frouin, V., Garavan, H., Gowland, P., Heinz, A., Ittermann, B., Martinot, J.-L., Martinot, M.-L. P., … Petrovic, P. (2020). Distinct brain structure and behavior related to ADHD and conduct disorder traits. Molecular Psychiatry, 25(11), 3020–3033. 10.1038/s41380-018-0202-6

Bender, A., Voytek, B., & Schaworonkow, N. (2023). Resting-state is not enough: Alpha and mu rhythms change shape across development, but lack diagnostic sensitivity (p. 2023.10.13.562301). bioRxiv. 10.1101/2023.10.13.562301

Benjamini, Y., & Hochberg, Y. (1995). Controlling the False Discovery Rate: A Practical and Powerful Approach to Multiple Testing. Journalof the Royal Statistical Society: Series B (Methodological), 57(1), 289–300. 10.1111/j.2517-6161.1995.tb02031.x

Biederman, J., Lanier, J., DiSalvo, M., Noyes, E., Fried, R., Woodworth, K. Y., Biederman, I., & Faraone, S. V. (2019). Clinical correlates of mind wandering in adults with ADHD. Journalof Psychiatric Research, 117, 15–23. 10.1016/j.jpsychires.2019.06.012

Boetzel, C., & Herrmann, C. S. (2021). Potential targets for the treatment of ADHD using transcranial electrical current stimulation. Progress in Brain Research, 264, 151–170. 10.1016/BS.PBR.2021.01.011

Boker, S., Neale, M., Maes, H., Wilde, M., Spiegel, M., Brick, T., Spies, J., Estabrook, R., Kenny, S., Bates, T., Mehta, P., & Fox, J. (2011). OpenMx: An Open Source Extended Structural Equation Modeling Framework. Psychometrika, 76(2), 306–317. 10.1007/s11336-010-9200-6

Brandeis, D., Loo, S. K., McLoughlin, G., Heinrich, H., & Banaschewski, T. (2018). Neurophysiology. In T. Banaschewski, D. Coghill, & A. Zuddas (Eds.), Oxford Textbook of Attention Deficit Hyperactivity Disorder (p. 0). Oxford University Press. 10.1093/med/9780198739258.003.0009

Bresnahan, S. M., Barry, R. J., Clarke, A. R., & Johnstone, S. J. (2006). Quantitative EEG analysis in dexamphetamine-responsive adults with attention-deficit/hyperactivity disorder. Psychiatry Research, 141(2), 151–159. 10.1016/j.psychres.2005.09.002

Cañigueral, R., Palmer, J., Ashwood, K. L., Azadi, B., Asherson, P., Bolton, P. F., McLoughlin, G., & Tye, C. (2022). Alpha oscillatory activity during attentional control in children with Autism Spectrum Disorder (ASD), Attention-Deficit/Hyperactivity Disorder (ADHD), and ASD+ADHD. Journalof Child Psychology and Psychiatry, 63(7), 745–761. 10.1111/jcpp.13514

Caplan, J. B., Bottomley, M., Kang, P., & Dixon, R. A. (2015). Distinguishing rhythmic from non-rhythmic brain activity during rest in healthy neurocognitive aging. NeuroImage, 112, 341–352. 10.1016/J.NEUROIMAGE.2015.03.001

Capp, S. J., Agnew-Blais, J., Lau-Zhu, A., Colvert, E., Tye, C., Aydin, Ü., Lautarescu, A., Ellis, C., Saunders, T., O’Brien, L., Ronald, A., Happé, F., & McLoughlin, G. (2023). Is quality of life related to high autistic traits, high ADHD traits and their Interaction? Evidence from a Young-Adult Community-Based twin sample. Journalof Autism and Developmental Disorders, 53(9), 3493–3508. 10.1007/s10803-022-05640-w

Cascade, E., Kalali, A. H., & Wigal, S. B. (2010). Real-World Data on: Attention Deficit Hyperactivity Disorder Medication Side Effects. Psychiatry (Edgmont (Pa.: Township)), 7(4), 13–15.

Chang, C.-Y., Hsu, S.-H., Pion-Tonachini, L., & Jung, T.-P. (2019). Evaluation of Artifact Subspace Reconstruction for Automatic Artifact Components Removal in Multi-channel EEG Recordings. IEEE Transactions on Biomedical Engineering, 1–1. 10.1109/tbme.2019.2930186

Chen, R., Liu, W., Wang, J.-J., Zhou, D.-D., & Wang, Y. (2024). Aperiodic components and aperiodic-adjusted alpha-band oscillations in children with ADHD. Journalof Psychiatric Research, 173, 225–231. 10.1016/j.jpsychires.2024.03.042

Compton, R. J., Gearinger, D., & Wild, H. (2019). The wandering mind oscillates: EEG alpha power is enhanced during moments of mind-wandering. Cognitive, Affective, & Behavioral Neuroscience, 1–8. 10.3758/s13415-019-00745-9

Cortese, S., Adamo, N., Giovane, C. D., Mohr-Jensen, C., Hayes, A. J., Carucci, S., Atkinson, L. Z., Tessari, L., Banaschewski, T., Coghill, D., Hollis, C., Simonoff, E., Zuddas, A., Barbui, C., Purgato, M., Steinhausen, H.-C., Shokraneh, F., Xia, J., & Cipriani, A. (2018). Comparative efficacy and tolerability of medications for attention-deficit hyperactivity disorder in children, adolescents, and adults: A systematic review and network meta-analysis. The Lancet Psychiatry, 5(9), 727–738. 10.1016/S2215-0366(18)30269-4

Crescenzo, F. D., Cortese, S., Adamo, N., & Janiri, L. (2017). Pharmacological and non-pharmacological treatment of adults with ADHD: A meta-review. BMJ Ment Health, 20(1), 4–11. 10.1136/eb-2016-102415

Dakwar-Kawar, O., Mentch-Lifshits, T., Hochman, S., Mairon, N., Cohen, R., Balasubramani, P., Mishra, J., Jordan, J., Cohen Kadosh, R., Berger, I., & Nahum, M. (2024). Aperiodic and periodic components of oscillatory brain activity in relation to cognition and symptoms in pediatric ADHD. Cerebral Cortex, 34(6), bhae236. 10.1093/cercor/bhae236

Deiber, M.-P., Hasler, R., Colin, J., Dayer, A., Aubry, J.-M., Baggio, S., Perroud, N., & Ros, T. (2020). Linking alpha oscillations, attention and inhibitory control in adult ADHD with EEG neurofeedback. NeuroImage: Clinical, 25, 102145. 10.1016/j.nicl.2019.102145

Delorme, A., & Makeig, S. (2004). EEGLAB: an open sorce toolbox for analysis of single -trail EEG dynamics including independent component anlaysis. Journalof Neuroscience Methods, 134, 9–21. 10.1016/j.jneumeth.2003.10.009

Diaz, B. A., Van Der Sluis, S., Moens, S., Benjamins, J. S., Migliorati, F., Stoffers, D., Den Braber, A., Poil, S.-S., Hardstone, R., Van’t Ent, D., Boomsma, D. I., De Geus, E., Mansvelder, H. D., Van Someren, E. J. W., & Linkenkaer-Hansen, K. (2013). The Amsterdam Resting-State Questionnaire reveals multiple phenotypes of resting-state cognition. Frontiers in Human Neuroscience, 7(JUL), 446. 10.3389/fnhum.2013.00446

Donoghue, T., Schaworonkow, N., & Voytek, B. (2021). Methodological considerations for studying neural oscillations. European Journalof Neuroscience. 10.1111/EJN.15361

Duan, W., Chen, X., Wang, Y.-J., Zhao, W., Yuan, H., & Lei, X. (2021). Reproducibility of power spectrum, functional connectivity and network construction in resting-state EEG. Journalof Neuroscience Methods, 348, 108985. 10.1016/j.jneumeth.2020.108985

Farokhzadi, F., Mohamadi, M. R., Khosli, A. K., Akbarfahimi, M., Beigi, N. A., & Torabi, P. (2020). Comparing the Effectiveness of the Transcranial Alternating Current Stimulation (TACS) and Ritalin on Symptoms of Attention Deficit Hyperactivity Disorder in 7-14-Year-Old Children. Acta Medica Iranica, 637–648. 10.18502/acta.v58i12.5156

Frohlich, F., & Riddle, J. (2021). Conducting double-blind placebo-controlled clinical trials of transcranial alternating current stimulation (tACS). Translational Psychiatry, 11(1), 1–12. 10.1038/s41398-021-01391-x

Günther, A., Hanganu-opatz, I. L., & Robinson, J. (2022). Neuronal oscillations: Early biomarkers of psychiatric disease *?* December, 1–9. 10.3389/fnbeh.2022.1038981

Hindriks, R., & van Putten, M. J. A. M. (2013). Thalamo-cortical mechanisms underlying changes in amplitude and frequency of human alpha oscillations. NeuroImage, 70, 150–163. 10.1016/j.neuroimage.2012.12.018

Klimesch, W. (1999). EEG alpha and theta oscillations reflect cognitive and memory performance: A rKlimesch, W. (1999). EEG alpha and theta oscillations reflect cognitive and memory performance: A review and analysis. Brain Research Reviews, 29(2-3), 169–195. Doi:10.1016/S016. *Brain Research Reviews*, *29*(2–3), 169–195. 10.1016/S0165-0173(98)00056-3

Klooster, D., Voetterl, H., Baeken, C., & Arns, M. (2024). Evaluating Robustness of Brain Stimulation Biomarkers for Depression: A Systematic Review of Magnetic Resonance Imaging and Electroencephalography Studies. Biological Psychiatry, 95(6), 553–563. 10.1016/j.biopsych.2023.09.009

Koehler, S., Lauer, P., Schreppel, T., Jacob, C., Heine, M., Boreatti-Hümmer, A., Fallgatter, A. J., & Herrmann, M. J. (2009). Increased EEG power density in alpha and theta bands in adult ADHD patients. Journalof Neural Transmission, 116(1), 97–104. 10.1007/s00702-008-0157-x

Kumral, D., Cesnaite, E., Beyer, F., Hofmann, S. M., Hensch, T., Sander, C., Hegerl, U., Haufe, S., Villringer, A., Witte, A. V., & Nikulin, V. V. (2022). Relationship between regional white matter hyperintensities and alpha oscillations in older adults. Neurobiology of Aging, 112, 1–11. 10.1016/j.neurobiolaging.2021.10.006

Lanier, J., Noyes, E., & Biederman, J. (2021). Mind Wandering (Internal Distractibility) in ADHD: A Literature Review. Journalof Attention Disorders, 25(6), 885–890. 10.1177/1087054719865781/ASSET/IMAGES/LARGE/10.1177_1087054719865781-FIG1.JPEG

Liu, Z.-X., Glizer, D., Tannock, R., & Woltering, S. (2016). EEG alpha power during maintenance of information in working memory in adults with ADHD and its plasticity due to working memory training: A randomized controlled trial. Clinical Neurophysiology, 127(2), 1307– 1320. 10.1016/j.clinph.2015.10.032

Loo, S. K., Hale, T. S., Macion, J., Hanada, G., McGough, J. J., McCracken, J. T., & Smalley, S. L. (2009). Cortical activity patterns in ADHD during arousal, activation and sustained attention. Neuropsychologia, 47(10), 2114–2119. 10.1016/j.neuropsychologia.2009.04.013

Loo, S. K., McGough, J. J., McCracken, J. T., & Smalley, S. L. (2018). Parsing heterogeneity in attention-deficit hyperactivity disorder using EEG-based subgroups. Journal of Child Psychology and Psychiatry, 59(3), 223–231. 10.1111/jcpp.12814

Malone, S. M., Burwell, S. J., Vaidyanathan, U., Miller, M. B., McGue, M., & Iacono, W. G. (2014). Heritability and Molecular-Genetic Basis of Resting EEG Activity: A Genome-Wide Association Study. Psychophysiology, 51(12), 1225–1245. 10.1111/psyp.12344

McCarthy, S., Wilton, L., Murray, M. L., Hodgkins, P., Asherson, P., & Wong, I. C. (2012). The epidemiology of pharmacologically treated attention deficit hyperactivity disorder (ADHD) in children, adolescents and adults in UK primary care. BMC Pediatrics, 12(1), 78. 10.1186/1471-2431-12-78

McLoughlin, G., Gyurkovics, M., & Aydin, Ü. (2022). What Has Been Learned from Using EEG Methods in Research of ADHD? In S. C. Stanford & E. Sciberras (Eds.), New Discoveries in the Behavioral Neuroscience of Attention-Deficit Hyperactivity Disorder (pp. 415–444). Springer International Publishing. 10.1007/7854_2022_344

McLoughlin, G., Makeig, S., & Tsuang, M. T. (2014). In search of biomarkers in psychiatry: EEG-based measures of brain function. American Journalof Medical Genetics Part B: Neuropsychiatric Genetics, 165(2), 111–121. 10.1002/ajmg.b.32208

McLoughlin, G., Palmer, J. A., Rijsdijk, F., & Makeig, S. (2014). Genetic Overlap between Evoked Frontocentral Theta-Band Phase Variability, Reaction Time Variability, and Attention-Deficit/Hyperactivity Disorder Symptoms in a Twin Study. Biological Psychiatry, 75(3), 238–247. 10.1016/J.BIOPSYCH.2013.07.020

Neale, M. C., & Maes, H. H. M. (2004). Methodology for Genetic Studies of Twins and Families. Kluwer Academic Publishers B.V. Dordrecht, The Netherlands.

Nimmo-Smith, V., Merwood, A., Hank, D., Brandling, J., Greenwood, R., Skinner, L., Law, S., Patel, V., & Rai, D. (2020). Non-pharmacological interventions for adult ADHD: A systematic review. Psychological Medicine, 50(4), 529–541. 10.1017/S0033291720000069

Ossadtchi, A., Shamaeva, T., Okorokova, E., Moiseeva, V., & Lebedev, M. A. (2017). Neurofeedback learning modifies the incidence rate of alpha spindles, but not their duration and amplitude. Scientific Reports, 7(1), 3772. 10.1038/s41598-017-04012-0

Pion-Tonachini, L., Kreutz-Delgado, K., & Makeig, S. (2019). ICLabel: An automated electroencephalographic independent component classifier, dataset, and website. NeuroImage, 198, 181–197. 10.1016/j.neuroimage.2019.05.026

Poil, S.-S., Bollmann, S., Ghisleni, C., O’Gorman, R. L., Klaver, P., Ball, J., Eich-Höchli, D., Brandeis, D., & Michels, L. (2014). Age dependent electroencephalographic changes in attention-deficit/hyperactivity disorder (ADHD). Clinical Neurophysiology, 125(8), 1626–1638. 10.1016/j.clinph.2013.12.118

Polanczyk, G., de Lima, M. S., Horta, B. L., Biederman, J., & Rohde, L. A. (2007). The Worldwide Prevalence of ADHD: A Systematic Review and Metaregression Analysis. American Journal of Psychiatry, 164(6), 942–948. 10.1176/ajp.2007.164.6.942

Ponomarev, V. A., Mueller, A., Candrian, G., Grin-Yatsenko, V. A., & Kropotov, J. D. (2014). Group Independent Component Analysis (gICA) and Current Source Density (CSD) in the study of EEG in ADHD adults. Clinical Neurophysiology, 125(1), 83–97. 10.1016/j.clinph.2013.06.015

Riddle, J., & Frohlich, F. (2021). Targeting neural oscillations with transcranial alternating current stimulation. Brain Research, 1765, 147491. 10.1016/j.brainres.2021.147491

Riddle, J., & Frohlich, F. (2023). Mental Activity as the Bridge between Neural Biomarkers and Symptoms of Psychiatric Illness. Clinical EEG and Neuroscience, 54(4), 399–408. 10.1177/15500594221112417

Rimfeld, K., Malanchini, M., Spargo, T., Spickernell, G., Selzam, S., McMillan, A., Dale, P. S., Eley, T. C., & Plomin, R. (2019). Twins Early Development Study: A Genetically Sensitive Investigation into Behavioral and Cognitive Development from Infancy to Emerging Adulthood. Twin Research and Human Genetics, 22(6), 508–513. 10.1017/thg.2019.56

Robertson, M. M., Furlong, S., Voytek, B., Donoghue, T., Boettiger, C. A., & Sheridan, M. A. (2019). EEG power spectral slope differs by ADHD status and stimulant medication exposure in early childhood. Journalof Neurophysiology, 122(6), 2427–2437. 10.1152/jn.00388.2019

Rodriguez-Larios, J., & Alaerts, K. (2020). EEG alpha-theta dynamics during mind wandering in the context of breath focus meditation: An experience sampling approach with novice meditation practitioners. *European Journalof Neuroscience*, May, 1–14. 10.1111/ejn.15073

Rodriguez-Larios, J., Bracho Montes de Oca, E. A., & Alaerts, K. (2021). The EEG spectral properties of meditation and mind wandering differ between experienced meditators and novices. NeuroImage, 245, 118669. 10.1016/J.NEUROIMAGE.2021.118669

Rodriguez-Larios, J., & Haegens, S. (2023). Genuine beta bursts in human working memory: Controlling for the influence of lower-frequency rhythms. Advances.in/Psychology, 1, 1–17. 10.56296/aip00006

Rodriguez-Larios, J., Wong, K. F., & Lim, J. (2024). Assessing the effects of an 8-week mindfulness training program on neural oscillations and self-reports during meditation practice. PLOS ONE, 19(6), e0299275. 10.1371/journal.pone.0299275

Schaworonkow, N., & Nikulin, V. V. (2019). Spatial neuronal synchronization and the waveform of oscillations: Implications for EEG and MEG. PLOS Computational Biology, 15(5), e1007055. 10.1371/journal.pcbi.1007055

Seli, P., Smallwood, J., Cheyne, J. A., & Smilek, D. (2015). On the relation of mind wandering and ADHD symptomatology. Psychonomic Bulletin & Review, 22(3), 629–636. 10.3758/s13423-014-0793-0

Skirrow, C., Mcloughlin, G., Banaschewski, T., Brandeis, D., Kuntsi, J., & Asherson, P. (2014). Normalisation of frontal theta activity following methylphenidate treatment in adult attention-deficit/hyperactivity disorder. European Neuropsychopharmacology, 25, 85–94. 10.1016/j.euroneuro.2014.09.015

Smit, C. M., Wright, M. J., Hansell, N. K., Geffen, G. M., & Martin, N. G. (2006). Genetic variation of individual alpha frequency (IAF) and alpha power in a large adolescent twin sample. International Journalof Psychophysiology, 61(2), 235–243. 10.1016/j.ijpsycho.2005.10.004

Soriano, J. R., Oca, E. B. M. de, Karaiskou, A.-I., Vuyst, H.-J. D., Rodriguez-Larios, J., Feng, N., Varon, C., & Alaerts, K. (2024). Bidirectional Alpha Power EEG Neurofeedback During a Focused Attention Meditation Practice in Novices. NeuroRegulation, 11(3), Article 3. 10.15540/nr.11.3.230

Taberna, G. A., Guarnieri, R., & Mantini, D. (2019). SPOT3D: Spatial positioning toolbox for head markers using 3D scans. Scientific Reports 2019 9:1, 9(1), 1–9. 10.1038/s41598-019-49256-0

Taberna, G. A., Samogin, J., Marino, M., & Mantini, D. (2021). Detection of resting-state functional connectivity from high-density electroencephalography data: Impact of head modeling strategies. Brain Sciences, 11(6), 741. 10.3390/BRAINSCI11060741/S1

Valdés-Hernández, P. A., Ojeda-González, A., Martínez-Montes, E., Lage-Castellanos, A., Virués-Alba, T., Valdés-Urrutia, L., & Valdes-Sosa, P. A. (2010). White matter architecture rather than cortical surface area correlates with the EEG alpha rhythm. NeuroImage, 49(3), 2328–2339. 10.1016/j.neuroimage.2009.10.030

van Beijsterveldt, C. E. M., & van Baal, G. C. M. (2002). Twin and family studies of the human electroencephalogram: A review and a meta-analysis. Biological Psychology, 61(1), 111–138. 10.1016/S0301-0511(02)00055-8

van Dongen-Boomsma, M., Lansbergen, M. M., Bekker, E. M., Sandra Kooij, J. J., van der Molen, M., Kenemans, J. L., & Buitelaar, J. K. (2010). Relation between resting EEG to cognitive performance and clinical symptoms in adults with attention-deficit/hyperactivity disorder. Neuroscience Letters, 469(1), 102–106. 10.1016/j.neulet.2009.11.053

Veniero, D., Vossen, A., Gross, J., & Thut, G. (2015). Lasting EEG/MEG Aftereffects of Rhythmic Transcranial Brain Stimulation: Level of Control Over Oscillatory Network Activity. Frontiers in Cellular Neuroscience, 9. 10.3389/fncel.2015.00477

Voetterl, H. T. S., Sack, A. T., Olbrich, S., Stuiver, S., Rouwhorst, R., Prentice, A., Pizzagalli, D. A., van der Vinne, N., van Waarde, J. A., Brunovsky, M., van Oostrom, I., Reitsma, B., Fekkes, J., van Dijk, H., & Arns, M. (2023). Alpha peak frequency-based Brainmarker-I as a method to stratify to pharmacotherapy and brain stimulation treatments in depression | Nature Mental Health. Nature Mental Health. 10.1038/s44220-023-00160-7

von Stein, A., & Sarnthein, J. (2000). Different frequencies for different scales of cortical integration: From local gamma to long range alpha/theta synchronization. International Journalof Psychophysiology, 38(3), 301–313. 10.1016/S0167-8760(00)00172-0

Vossen, A., Gross, J., & Thut, G. (2015). Alpha Power Increase After Transcranial Alternating Current Stimulation at Alpha Frequency (α-tACS) Reflects Plastic Changes Rather Than Entrainment. Brain Stimulation, 8(3), 499–508. 10.1016/J.BRS.2014.12.004

Woltering, S., Jung, J., Liu, Z., & Tannock, R. (2012). Resting state EEG oscillatory power differences in ADHD college students and their peers. Behavioral and Brain Functions, 8(1), 60. 10.1186/1744-9081-8-60

